# Short-term mechanisms, long-term consequences: molecular effects of ocean acidification on juvenile snow crab

**DOI:** 10.64898/2026.02.04.703865

**Authors:** Laura H Spencer, Ingrid B. Spies, Jennifer L. Gardner, Steven B. Roberts, W. Christopher Long

## Abstract

Understanding how marine species tolerate acidified conditions is critical for predicting biological responses to ocean change. A recent one-year experiment (Long 2026) found that juvenile snow crab (*Chionoecetes opilio*) maintain growth and molting under acidification (pH 7.8, 7.5), and survival begins to decline only after ∼250 days under severe acidification (pH 7.5). In this companion study, we characterized whole-transcriptome responses after 8 hours and 88 days of exposure to identify molecular mechanisms underlying short-term tolerance and chronic effects of ocean acidification. The immediate transcriptional response involved strong activation of genes associated with mitochondrial metabolism and biogenesis, protein homeostasis, cuticle maintenance, and immune modulation, processes shared between moderate and severe treatments but of greater magnitude under severe acidification. After 88 days, expression patterns diverged, revealing sustained upregulation of stress- and damage-mitigation pathways in the severe treatment (pH 7.5) compared to the moderate treatment (pH 7.8). These findings indicate that crabs in severe acidification are likely to experience chronic OA stress that precedes outward physiological effects, and provides a mechanistic basis for delayed mortality. We further highlight potential early indicators of chronic acidification stress in snow crab, among which a gene likely coding for carbonic anhydrase 7 (CA7, GWK47_031192) appears to be the most promising biomarker.

**Summary Statement:** Juvenile snow crabs tolerate ocean acidification through flexible gene expression, but prolonged exposure reveals hidden cellular stress that helps explain delayed mortality.

## 1. Introduction

Atmospheric and ocean conditions are changing at an unprecedented rate compared to the geologic record. Since monitoring of atmospheric carbon dioxide began at Mauna Loa Research Station (Keeling et al., 1976), concentrations have risen by ∼33%, from 316 ppm in 1959 to 425 ppm in 2024 (based on annual means, https://gml.noaa.gov/ccgg/trends/data.html). Roughly 25% of this additional CO₂ has been absorbed by the ocean (DeVries, 2022), driving a decrease in ocean pH of 0.1 units over the past 50 years (DeVries, 2022). Because pH is on a logarithmic scale, this represents an approximately 25% increase in proton concentration ([H⁺]), accompanied by reductions in the saturation states of dissolved calcium carbonates (Doney et al., 2009). Reduced carbonate availability can impair shell and skeletal formation in calcareous marine organisms by impeding biomineralization (Figuerola et al., 2021). Some organisms cannot fully regulate internal acid-base balance (Auzoux-Bordenave et al., 2021), while others maintain internal pH levels through buffering (Fehsenfeld & Weihrauch, 2017), but at an energetic cost (Meseck et al., 2016). Consequently, the observed shifts in the ocean carbon cycle, collectively termed ocean acidification (Caldeira & Wickett, 2003), pose significant risks to marine communities by altering the physiology and productivity of calcareous species.

Large benthic decapod crabs are calcareous species that play critical ecological and economic roles as predators, prey, and major fisheries targets (Boudreau & Worm, 2012). Many of these crabs may be especially vulnerable to ocean acidification, as they live in high-latitude regions that are acidifying (and warming) more rapidly than lower latitudes (Fabry et al., 2009). Decapod crabs, like other calcareous taxa, are typically most sensitive to changes in pH (commonly used as a proxy for acidification) during early life stages, as embryos, larvae, and juveniles (Long et al., 2013; Przeslawski et al., 2015). However, as with many marine taxa (Kroeker et al., 2010; Vargas et al., 2017), decapod responses to ocean acidification are notoriously diverse, differing among species, among populations within species, and even among individuals within populations (Bednaršek et al., 2021; McElhany & Busch, 2024; Siegel et al., 2022). Understanding the causes of this variation is now a central focus. After decades of field observations and laboratory experiments, research efforts are turning toward the development of mechanistic models to distinguish tolerant versus sensitive groups (Leung et al., 2022), with the goal of improving management and conservation strategies.

The diversity of crab species that co-occur in the North Pacific Ocean provides opportunities to identify factors that differentiate OA-tolerant crabs from those that are OA-sensitive (Knauber et al., 2023; Otto & Jamieson, 2001). Experimental work has examined the responses of king (red, blue, golden), snow, tanner, and dungeness crabs to acidification, all of which are found in Alaskan waters (Kruse et al., 2025). Across these species, early life stages often experience reduced survival, slower growth, or impaired calcification, although the severity of effects varies (Coffey et al., 2017; Long et al., 2013, 2016, 2019, 2021, 2024; Miller et al., 2016; Swiney et al., 2016; Trigg et al., 2019). Juvenile red king crab (*Paralithodes camtschaticus*) are among the most sensitive, with 100% mortality reported at pH 7.5 within 95 days (Long et al., 2013; Swiney et al., 2017, although see Long et al. 2024 and Spencer et al. 2024), and this sensitivity is thought to contribute to recent declines in productivity in the southeastern Bering Sea (Litzow et al., 2025). Tanner crab (*Chionoecetes bairdi)* are similarly vulnerable, as adults exhibit impaired reproductive output when held at low pH (Swiney et al., 2016), larvae show significantly reduced starvation-survival under acidified conditions (Long et al., 2016), and long-term juvenile exposures result in reduced calcification, slower growth, and increased mortality at pH 7.5 (Long et al., 2013).

Snow crab (*Chionoecetes opilio*) stand out as a striking exception. Despite being closely related to Tanner crab and even capable of hybridizing with them, snow crab appear relatively tolerant to OA. Embryos and larvae consistently show little to no direct sensitivity to acidification, with embryo morphology, hatching success, and larval condition largely unaffected even at the most acidified treatment (pH 7.5, Long et al., 2023). Adult calcification is similarly unaffected (Algayer et al., 2023). Long (2026) recently demonstrated that this tolerance largely extends to the juvenile stage, as juvenile morphology and molting were unaffected after a full year of exposure to moderate (pH 7.8) and severe (pH 7.5) acidification. However, elevated mortality began after 250 days in the severe treatment and ultimately resulted in ∼40% higher mortality by the end of the experiment. This long-term decline highlights that even apparently resilient stages can experience cumulative physiological costs over time, and that chronic exposures, more akin to real-world ocean conditions, may reveal impacts that short-term observations overlook.

This study advances understanding of both the short-term acclimation mechanisms and the long-term consequences of ocean acidification in snow crab. Using individuals from Long (2026), we applied transcriptomics to assess gene expression profiles after short-term (8 h) and intermediate (88 d) exposure in the moderate and severe OA treatments. Transcriptomics captures the complete set of expressed genes, providing a comprehensive view of active physiological pathways and their energetic demands (Page & Lawley, 2022). Long (2026) observed that mortality under severe OA began after ∼250 days, while our data capture much earlier time points. These snapshots provide a unique opportunity to address three key questions: (1) How do snow crabs acclimate to acidification in the short term? (2) Do these acclimation mechanisms persist over longer exposures? and (3) Which molecular pathways differ between crabs exposed to severe (pH 7.5) versus moderate (pH 7.8) acidification, that is, which mechanisms ultimately drive mortality under chronic acidification? Finally, we identify candidate biomarkers that could be used to determine whether snow crabs and related species (e.g., Tanner crabs and hybrids) are experiencing chronic OA stress.

## 2. Methods

### 2.1 Experimental Design and Sampling

This work was conducted in concert with a larger study on how ocean acidification affects snow crab (*Chionoecetes opilio*); full methodological details are available in Long (2026); crabs in this study were held in the same tanks, under the same treatment conditions, and fed the same diet. Briefly, snow crab juveniles (wet weight: 0.24 ± 0.03 g; carapace width: 8.3 ± 0.3 mm) were collected by trawl from the Bering Sea in April 2021 and transported to the Kodiak Laboratory, where they were acclimated in flow-through seawater for 2 weeks (4°C, ambient salinity), then randomly assigned to one of three pH treatments: Control / ambient pH_T_ (∼8.0), moderate acidification (pH 7.8), or severe acidification (pH 7.5). These levels are relevant based on snow crab distribution in the Bering Sea where seasonal lows of pH around 7.5 occur (Mathis et al., 2014), and align with previous experiments (Long et al., 2013; Spencer et al., 2024). Crabs were housed individually in mesh-bottom PVC inserts in 380 L tanks chilled to 4°C, and target pH was maintained by CO_2_ bubbling with Durafet III probe feedback. pH was measured three times a week with a Durafet III probe and the salinity weekly with Mettler Toledo InLab 731-2m salinity probe. Alkalinity was calculated from weekly salinity values using the established salinity-alkalinity relationship for the Gulf of Alaska (Evans et al. 2015) and the other carbonate chemistry parameters were calculated with the seacarb package in R (R 4.2.3, Vienna, Austria) (Table 1). Temperature was maintained within a 1°C range suitable for rearing snow crab (2.7 – 3.7°C).

**Table 1.** Summary of water chemistry parameters, temperature, and RNA-Seq sampling scheme. Parameter values represent mean ± SD calculated from repeated measurements taken from the start of the experiment (Day 1; April 23, 2021) through each corresponding sampling date. Salinity and alkalinity values without associated error indicate parameters measured on Day 1 only. Temperature and pH were measured three times per week, salinity every week, and all other parameters were calculated from these measurements.

|  | Day 1 |  |  | Day 88 |  |
| --- | --- | --- | --- | --- | --- |
| OA Treatment | Ambient<br>(pH ~8.0) | Moderate OA<br>(pH ~7.8) | Severe OA<br>(pH ~7.5) | Moderate OA<br>(pH ~7.8) | Severe OA<br>(pH ~7.5) |
| Tissue Sample Time | 0 hrs | 8 hrs | 8 hrs | 88 days | 88 days |
| Number of RNA-Seq libraries<br>(individual whole-body crabs) | 12 | 10 | 12 | 13 | 14 |
| Temperature | 3.2 ± 0.1 | 2.8 ± 0.0 | 2.8 ± 0.2 | 3.61 ± 0.4 | 3.69 ± 0.5 |
| Salinity | 32.1 | 32.1 | 32.0 | 31.5 ± 0.4 | 31.6 ± 0.5 |
| pH<sub>T</sub> | 7.96 ± 0.01 | 7.76 ± 0.01 | 7.52 ± 0.01 | 7.79 ± 0.02 | 7.49 ± 0.04 |
| Alkalinity (mmol/kg) | 2.17 | 2.17 | 2.17 | 2.14 ± 0.02 | 2.15 ± 0.02 |
| pCO<sub>2</sub> (μatm) | 463 ± 7 | 764 ± 19 | 1,348 ± 19 | 711 ± 44 | 1,459 ± 132 |
| HCO<sub>3</sub><sup>-</sup> (mmol/kg) | 1.972 ± 0.002 | 2.045 ± 0.002 | 2.092 ± 0.001 | 2.005 ± 0.020 | 2.074 ± 0.021 |
| CO<sub>3</sub><sup>-2</sup> (mmol/kg) | 0.079 ± 0.001 | 0.050 ± 0.001 | 0.030 ± 0.001 | 0.054 ± 0.003 | 0.028 ± 0.003 |
| DIC (mmol/kg) | 2.08 ± 0.001 | 2.14 ± 0.002 | 2.20 ± 0.002 | 2.10 ± 0.02 | 2.18 ± 0.02 |
| $\Omega_{\text{Aragonite}}$ | 1.20 ± 0.01 | 0.77 ± 0.01 | 0.45 ± 0.01 | 0.82 ± 0.05 | 0.43 ± 0.04 |
| $\Omega_{\text{Calcite}}$ | 1.91 ± 0.02 | 1.23 ± 0.02 | 0.72 ± 0.01 | 1.3 ± 0.07 | 0.69 ± 0.06 |

Tissue samples were collected for gene expression analysis on days 1 and 88 of treatment, and included four treatments: (1) a control treatment using crabs held at ambient conditions, after 8 hours of exposure to (2) pH 7.8 and (3) and pH 7.5, and after 88 days of exposure to (4) pH 7.8 and (5) pH 7.5 (Figure 1, Table 1). Prior to sampling, all crabs were starved for 24 hrs, then sacrificed through humane euthanasia by puncturing the carapace through the cardiac region, and preserved directly in RNA later, which was then held overnight at 4°C prior to being transferred to -80°C. Target sample size was 12 crabs per treatment and more crabs were held over the 88 day exposure than needed to account for potential mortality.

**Figure 1.**
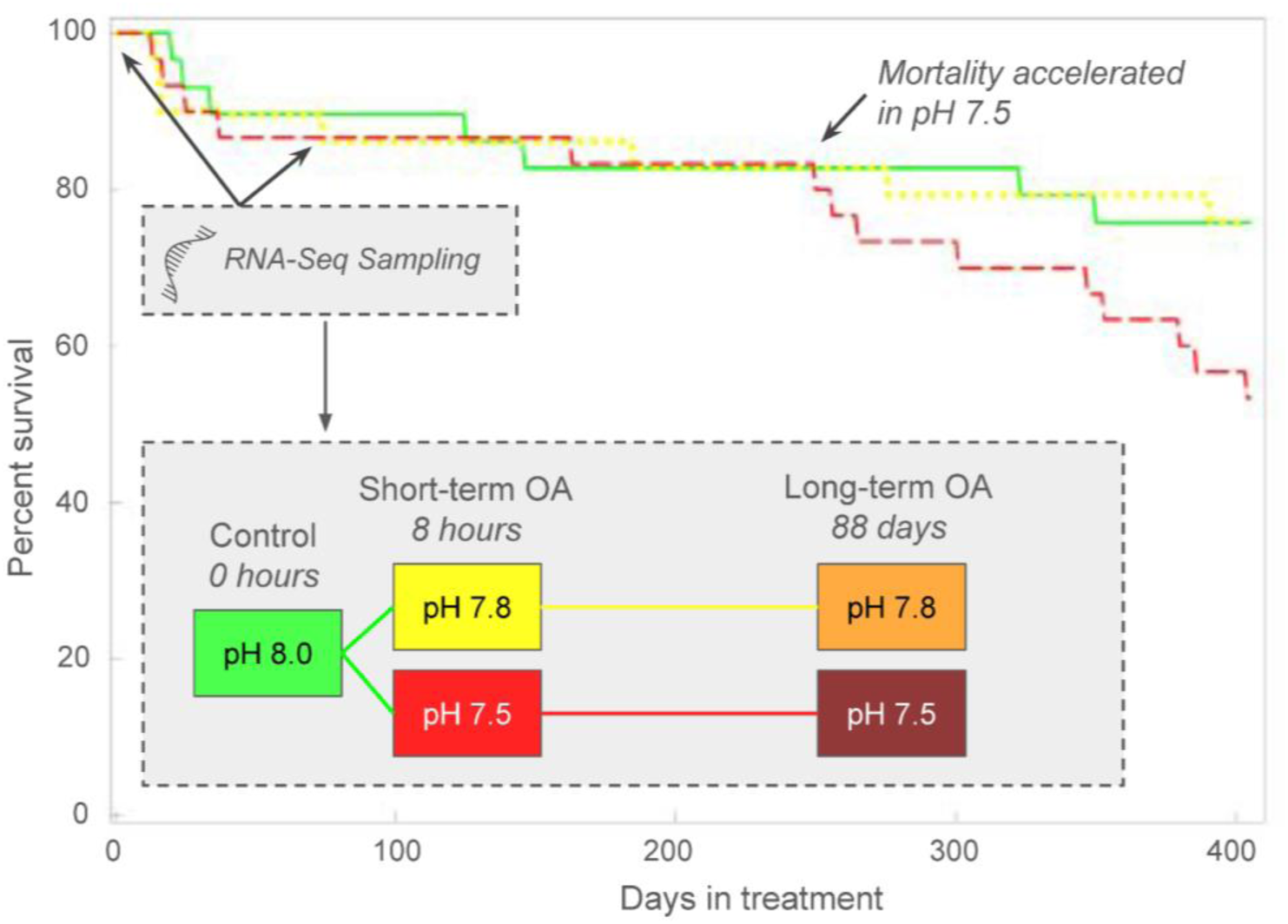
Juvenile snow crab survival did not vary among treatments until day 250, when higher mortality began to occur in the lowest pH treatment (pH 7.5). Tissues were sampled for gene expression on Day 1 from control treatment (0 hrs) and two OA treatments (8 hrs) to examine short-term acclimation mechanisms, and on Day 88 (two OA treatments) to examine long-term consequences of OA. Growth, intermolt duration, and morphology was unaffected by a 396-day exposure to varying pH treatments (Long 2026). Figure adapted from Long (2026).

### 2.2 RNASeq data generating and processing

Total RNA was isolated from RNAlater-preserved and homogenized whole-body crab tissue. Specimens were transferred from RNAlater into individual tubes containing cold TRIzol® reagent (500 µL) (Invitrogen, Carlsbad, CA, USA) and ∼five scoops of silica beads, then homogenized using a SPEX SamplePrep Geno/Grinder®. RNA was extracted following the manufacturer’s TRIzol® protocol. RNA-seq libraries were prepared using the Universal Plus™ mRNA-Seq with NuQuant kit (Tecan Genomics, Redwood City, CA, USA) for Illumina sequencing. Paired-end sequencing (150 bp) was performed on two lanes of an Illumina NovaSeq 6000 Sequencing System (Illumina, Inc., San Diego, CA, USA), with each sample split across both lanes. Raw sequence data were demultiplexed, concatenated by sample, and trimmed using Cutadapt v4.5 (Martin 2011) to remove Illumina adapters, flanking Ns, reads <50 bp, and low-quality ends (minimum quality scores of 20 and 15 for the 5′ and 3′ ends, respectively). Raw and trimmed reads were assessed using FastQC (Andrews, 2010) and MultiQC (Ewels et al., 2016).

Trimmed paired-end RNA-seq reads from 5 treatments (Table 1) were aligned to the *C. opilio* genome assembly from NCBI (accession: GCA_016584305.1_ASM1658430v1) using STAR v2.7.10a in splice-aware single-pass mode, with the options --quantMode GeneCounts and --outFilterMultimapNmax 30. Genome indexing was performed with the corresponding GTF annotation file to incorporate known splice junctions, using sjdbOverhang=149 to match the 150 bp read length. Gene counts, i.e. the number of paired-end fragments mapping to each gene, were extracted from STAR’s ReadsPerGene.out.tab files. One low-depth outlier sample (“S45”, ∼680K fragments) from the 88-day moderate OA treatment and another outlier from the 8-hr moderate OA treatment which influenced differential expression analysis (“S60”, 3.4M fragments) were excluded from analyses, leaving 10-14 samples per treatment (Table 1). Genes were filtered to remove those that were expressed at low levels, i.e. genes with fewer than 10 fragments in ≥10% of samples. Of the 6.13B trimmed read pairs, a total of 4.84B aligned to the snow crab genome (mean alignment rate of 78.9 ±3.5%), 2.16B of which mapped to annotated gene features. This resulted in appr 35% of raw reads and 45% of aligned reads being represented in the final gene count matrix, which included data for 11,097 unique genes to be analyzed for differential expression analysis.

Gene function was determined using a combination of available NCBI annotations and DIAMOND v2.1.8 blastx (Buchfink et al., 2015) searches against the UniProt/Swiss-Prot protein database (Release 2025_03)(UniProt Consortium, 2021), using an e-value cutoff of 1e-10 and retaining only the top hit. Header metadata from the snow crab coding sequence (cds) FASTA file was parsed to extract gene ids and functional tokens (e.g., GO terms, InterPro, Pfam). These were linked to external GO annotations using mapping files (interpro2go, pfam2go) from the Gene Ontology Consortium. For genes lacking functional labels, top DIAMOND matches were used to assign likely gene names and descriptions. Gene-to-GO mappings were compiled for downstream enrichment analyses.

### 2.3 Expression Analysis

Gene expression analyses were performed in R v4.3.2 using RStudio interface V2023.09.1+494 (R Core Team 2021; RStudio Team 2020). Unless otherwise specified, α = 0.05 and error bars represent ±1 SD.

#### 2.3.1 Global exploration and latent heterogeneity

We first performed an exploratory PCA of all analyzed genes with DESeq2’s plotPCA() on variance-stabilized counts to assess global expression patterns. This revealed latent, unknown sources of variation not aligned with known library prep, alignment, or biological variables in PC1-PC3 (Figure S1). To control for this latent heterogeneity, we used surrogate variable analysis (SVA) with the *sva* R package (Leek et al. 2012). Specifically, we preserved the pH treatment effect (∼ treatment) when estimating SVs, computed the number of SVs with num.sv(), and estimated them with svaseq() on variance-stabilized counts. The SVs were regressed from the VST matrix using limma::removeBatchEffect, and we then performed PCA again with stats::prcomp on the SV-adjusted matrix.

Global differences among treatments were assessed with permutational MANOVA (PERMANOVA) on the SV-adjusted VST matrix using adonis2 from the vegan R package (Oksanen et al. 2022). Pairwise treatment differences were evaluated with pairwise.adonis (pairwiseAdonis package, Martinez 2017). To identify genes contributing most strongly to the observed treatment differences we extracted the top 1% of genes with the largest positive and negative loadings on PC1 (0.5% per sign), a threshold selected based on the distribution of loadings (Figure S2). PCA biplot (PC1xPC2) visualized global expression variability, alongside tile plots which visualized expression patterns of the top 1% of genes contributing to each positive (n=55) and negative (n=55) ends of PC axes. Inter-individual variability within OA treatments was assessed by calculating Euclidean distances of samples to their treatment centroids in PCA space (PC1 x PC2). Distances to centroid were log-transformed and compared among treatments using ANOVA with Tukey’s HSD post hoc tests.

#### 2.3.2 Differential expression analysis

To identify genes that varied among treatments (differentially expressed genes, or DEGs), we analyzed raw counts with DESeq2 (default settings) (Costa-Silva et al., 2017; Love et al., 2014).

As recommended (Jaffe et al., 2015), we did not use the SV-regressed matrix for hypothesis tests, and instead all automatically detected SVs (n=7) from svaseq() were included as covariates in the DESeq2 design together with treatment (∼ SV1 + SV2 + … + SV7 + treatment). We then fit the model with DESeq(), and significant DEGs were defined by FDR-adjusted p-value (P_adj_ < 0.05, hereafter FDR) among treatment contrasts. We compared differentially expressed gene (DEG) lists to evaluate:

1. Short-term acclimation to moderate OA: pH 7.8 at 8 h vs. control at 0 h
2. Short-term acclimation to severe OA: pH 7.5 at 8 h vs. control at 0 h
3. Long-term consequences of severe OA: pH 7.5 vs. pH 7.8 at 88 days.

To identify potential gene targets for biomarker development of ocean acidification (OA) tolerance and stress, we also examined expression patterns across short-term (8 h, 88 d) and long-term exposures relative to time-0 controls. Genes were prioritized if they showed (1) consistently elevated expression under both short-term OA treatments and long-term exposure compared to control, or (2) differential expression among chronic OA treatments that also differed from time-0 controls.

For all differential expression analyses, outliers were handled using *DESeq2*’s built-in replacement method, where Cook’s Distance was used to identify influential outliers and then the original count values were replaced with trimmed means. To ensure that no outliers were influencing our DEG lists, we also used an iterative Leave-One-Out (iLOO) approach to identify and remove DEGs with outlier samples (n=114 genes) (George et al., 2015).

### 2.4 Functional Analyses

Differentially expressed gene sets were characterized by identifying enriched biological processes based on Gene Ontology (GO) terms and Uniprot keywords. For each set of differentially expressed genes, two enrichment analyses were performed to determine the functions of genes with higher and lower expression in OA treatments compared to control, or in severe OA compared to moderate OA. Enrichment was performed using Uniprot species IDs and the DAVID Bioinformatics Resources (v2023). For each enrichment analysis, the full set of annotated expressed genes was used as the background, P-values were adjusted using the false discovery rate (“FDR”) method, and enriched terms were defined as those with P_adj_ < 0.1 and at least two contributing genes. In cases where enriched GO terms and Uniprot keywords were the same, we only report GO terms.

All bioinformatics analyses were conducted on the NOAA Fisheries high-performance computing (HPC) cluster “Sedna”, maintained by the Northwest Fisheries Science Center (NWFSC) in Seattle, Washington. Code and data, including gene count files produced by STAR and augmented annotation files are available on GitHub (Spencer et al. 2026).

## 3. Results

### 3.1 Summary of growth, morphometrics, and survival

Full results are reported in Long (2026) and summarized here. Across treatments, crabs molted up to three times during the experiment, with most completing their first molt within one month and prior to the Day 88 sampling. Growth metrics (wet mass, carapace width, intermolt duration, and morphometric traits) did not differ among pH treatments. Survival rates through the 88-day gene expression sampling point did not differ by treatment; however, after continued monitoring through 396 days, survival diverged by treatment (Figure 1). Differential mortality began after day 250, resulting in >75% survival in ambient and moderate OA treatments but 50% survival under severe OA, where mortality was often associated with molting.

### 3.2 Global expression variability

We analyzed 2.1B read pairs from 62 samples which mapped to a total of 11,097 putative genes (per-sample average of 34.9 ± 7.0M read pairs mapped to 11,000 ± 44 genes; overall mean mapping rate was 78.9 ± 3.5%). After controlling for unknown surrogate variables (see methods & Figure S1), principal component analysis using all genes differentiated individuals by treatment and time along PC1 (9.1% variability) and PC2 (5.7% variability) (Figure 2), and permANOVA detected significant differences in multivariate space among treatments (F_4,56_=2.55, P=1.0e^−3^).

**Figure 2.**
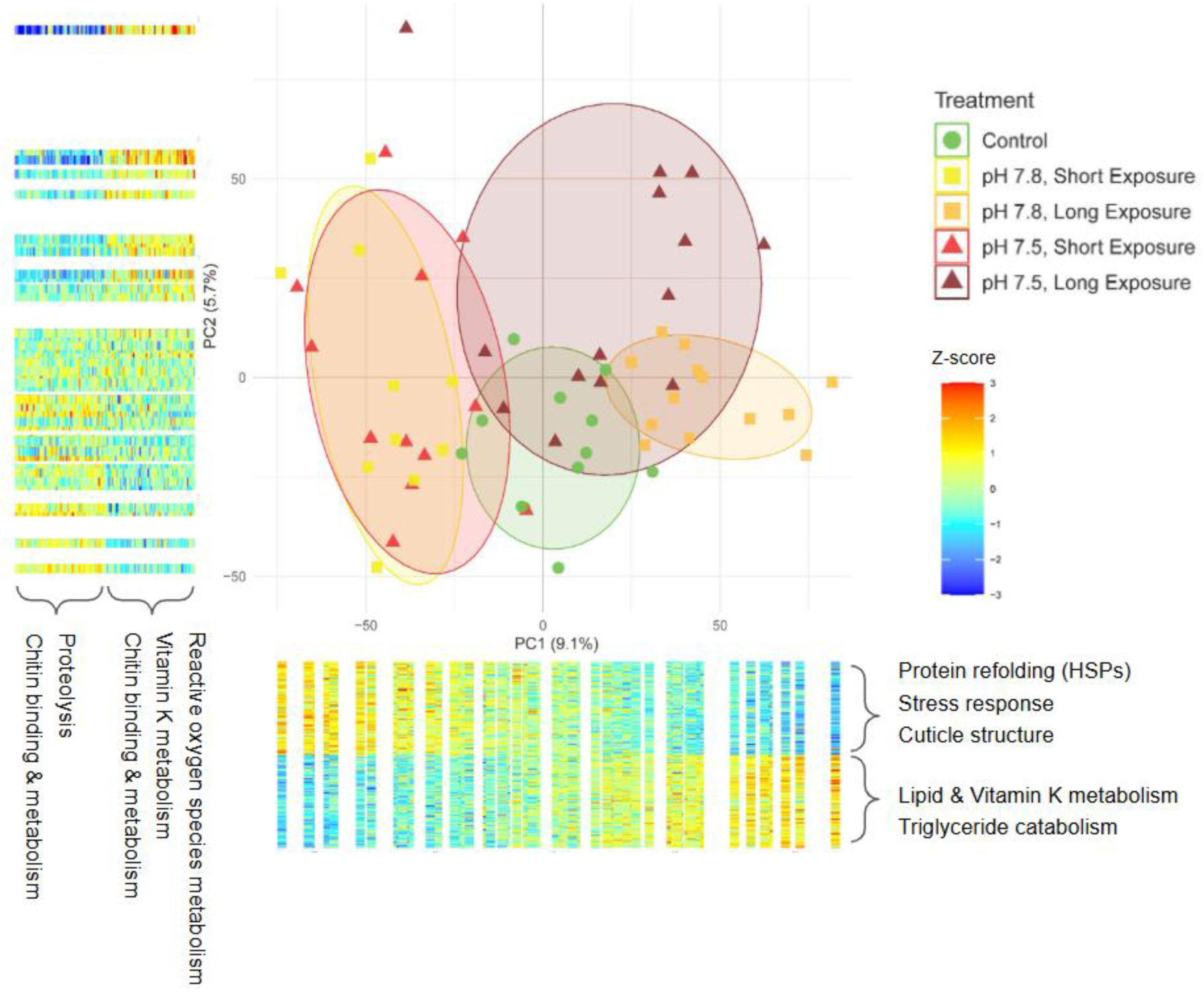
Principal component analysis biplot constructed from all analyzed transcripts, showing distinct clustering by pH treatment and exposure duration. PCA was performed on variance-stabilized gene expression data after regressing out latent variation with surrogate variable analysis (SVA). Adjacent heatmaps display z-transformed expression values for the top 1% of genes (n = 110, red = highest expression, blue = lowest expression) with the largest loadings on PC1 and PC2, with enriched processes shown for each gene set. For PC1, rows represent genes and columns represent individual crabs ordered by PC1 score; for PC2, columns represent genes and rows represent individual crabs ordered by PC2 score.

The genes with the strongest PC1 and PC2 loadings represent those most variable among the ∼11,097 genes analyzed and shaped the overall PCA clustering. For each axis, we performed enrichment analysis on the top 1% of genes, defined as the 0.5% with the most positive loadings and the 0.5% with the most negative loadings.

Genes with the largest negative PC1 loadings were those most highly expressed in the two short-term OA treatments (red and yellow in Figure 2), were strongly enriched for the Uniprot keyword stress response (FDR=2e^−9^) and GO processes protein refolding (FDR=5e^−11^), chaperone cofactor-dependent protein refolding (FDR=2e^−8^), and cellular response to unfolded protein (4.3e^−2^), due to high prevalence of molecular chaperones (e.g., HSP90). The ten genes with the largest negative PC1 loadings included two heat shock proteins (HSP90AA1, HSP71_1), two cuticle proteins (CUPA3_0, CUPA3_2), a detoxicant (UGT2B15), and five hypothetical proteins (Table S2). Genes with the largest positive PC1 loadings had higher expression in the long-term OA treatments (orange and dark red) and were enriched for GO processes vitamin K metabolic process (FDR=0.0027), triglyceride catabolic process (FDR=0.0028), and lipid metabolic process (FDR=0.0028). The ten genes with the largest positive PC1 loadings included two pancreatic lipase-related proteins (PNLIPRP2_1, PNLIPRP1_0), three solute carrier transporters (SLC18B1_0, SLC15A1, MFSD12_1), an ankyrin structural protein (ANK1), a leukocyte receptor cluster member (LENG9), and three hypothetical proteins (Table S2).

Genes with the largest positive PC2 loadings had higher expression in many crab exposed to long-term severe OA and some crab exposed to short-term OA, were enriched for positive regulation of reactive oxygen species metabolic process (FDR=0.047) and vitamin K metabolic process (FDR=0.01), and the ten genes with highest PC2 loadings included an immune-related receptor (LENG9), a redox regulator (DOXA-1), a carboxylesterase (EST6_2), a carnitine biosynthesis enzyme (BBOX1_6), and six hypothetical proteins (Table S2). Genes with the largest negative PC2 loadings were those with highest expression in time-0 controls and some short-term OA individuals, were enriched for peptide metabolic process (FDR=0.038) and protolysis (FDR=0.070), and the top ten genes included a serine protease (PRSS55), an integumentary mucin (MUCB1), an antimicrobial peptide (ALF_2), a carboxypeptidase (CPB1), and six hypothetical proteins (Table S2). Finally, chitin binding proteins were prevalent in the top PC2 loadings, both positive and negative.

Inter-individual transcriptional variability, as defined in PC space, differed among OA treatments (F_4,56_=3.8, P=0.0083). Mean distances to each treatment centroid was 6.2 ± 2.5 in time-0 control, 9.2 ± 5.4 in moderate short-term OA, and 5.5 ± 2.6 in moderate long-term OA. In contrast, variability was highest under severe OA, with distances of 10.1 ± 4.4 (short-term) and 11.0 ± 5.5 (long-term), indicating increased expression heterogeneity under high acidification stress.

### 3.3 Short term acclimation mechanisms: differential expression at 8 hours

We compared gene expression of crab sampled from ambient pH at time-0 to those exposed to moderate (pH 7.8) and severe (pH 7.5) acidification for 8 h to identify short-term acclimation mechanisms.

In crabs held for 8-hr in moderate acidification (pH 7.8), 148 genes were differentially expressed (DEGs). Genes with higher expression patterns (n=61) in moderate OA were enriched for the Uniprot keywords respiratory chain (FDR=0.022) and electron transport (FDR=0.088), and annotated genes with large effect size (Log_2_FC>1) that were uniquely expressed in moderate OA included mandibular organ-inhibiting hormone (MOIH), solute carrier family 22 member 13 (SLC22A13), and dual oxidase maturation factor 1 (doxa-1) (Figure 3).

**Figure 3.**
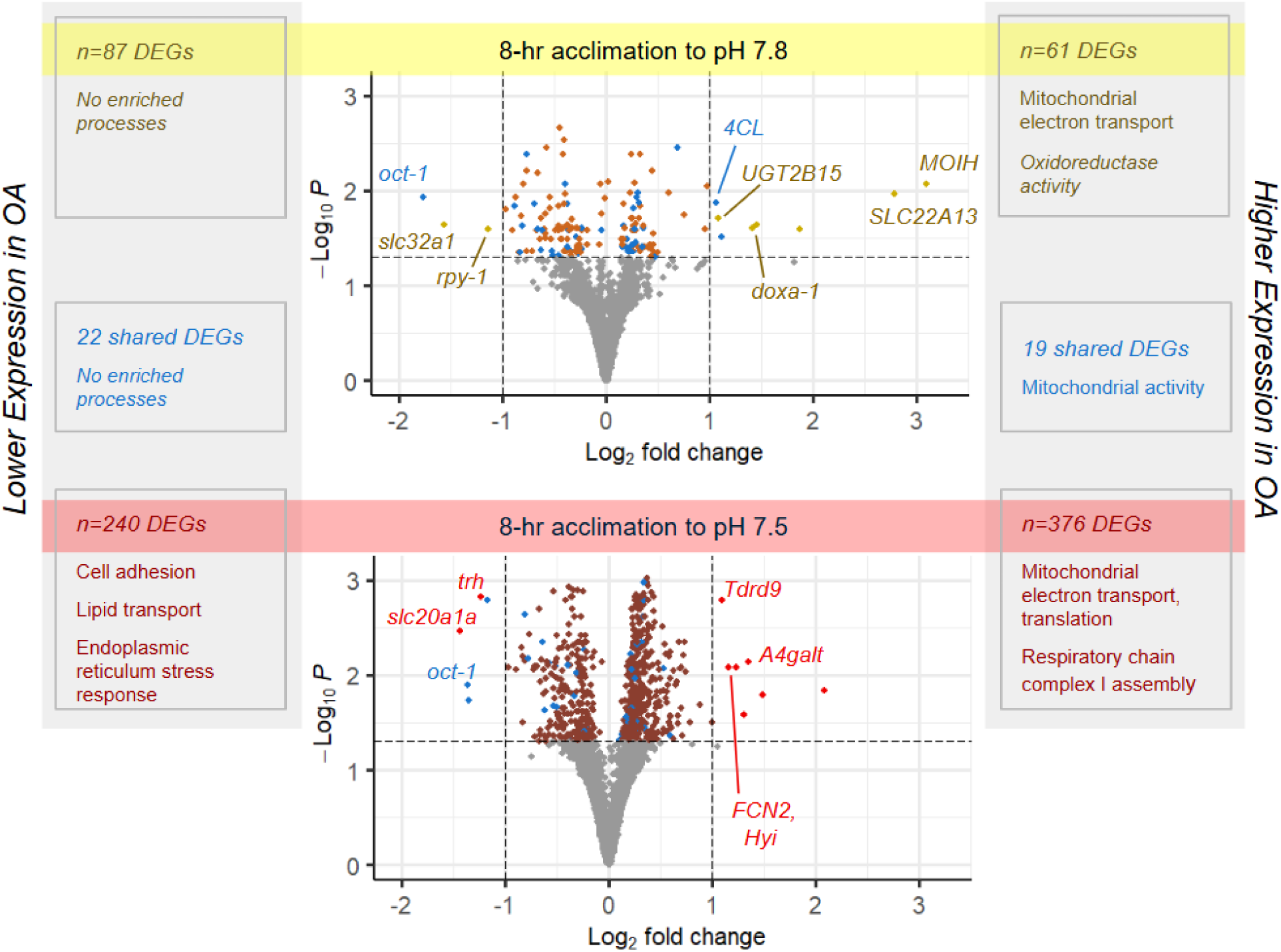
Differential gene expression and enriched biological processes in juvenile snow crab after 8-hr exposure to two acidification levels (pH 7.8, pH 7.5). Volcano plots show differentially expressed genes (DEGs) among time-0 control (pH 8.0) and OA treatments, which reveal a more robust transcriptional response in the more severe OA treatment (pH 7.5), and highlight annotated genes with >50% fold difference (|L_2_FC|>0.58). Panels show the number of DEGs in each 8-hr OA treatment by expression profile (lower, higher), those shared across both OA treatments (in blue, see Table 2), and associated biological processes (enriched Gene Ontology terms and Uniprot Keywords).

**Table 2.**
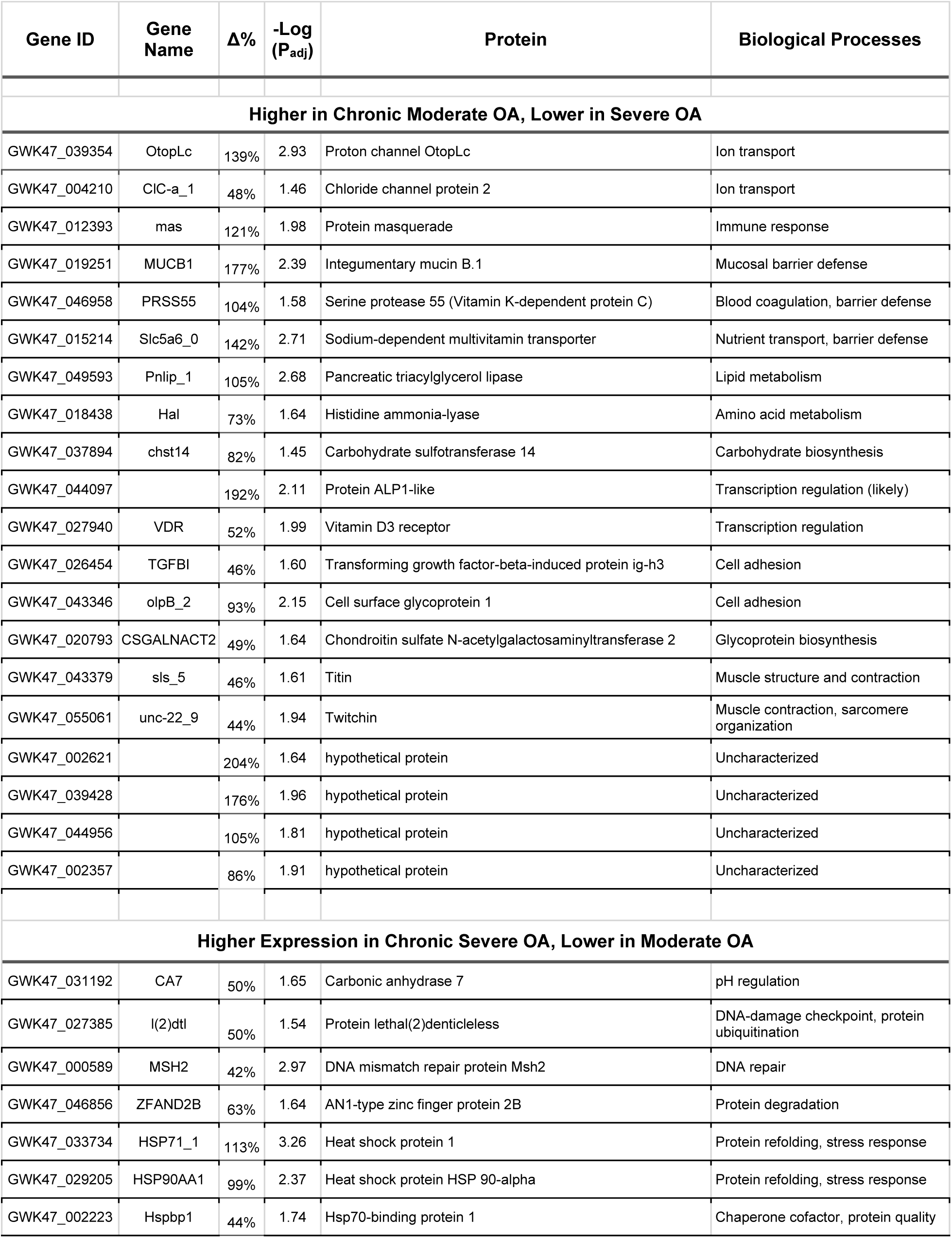
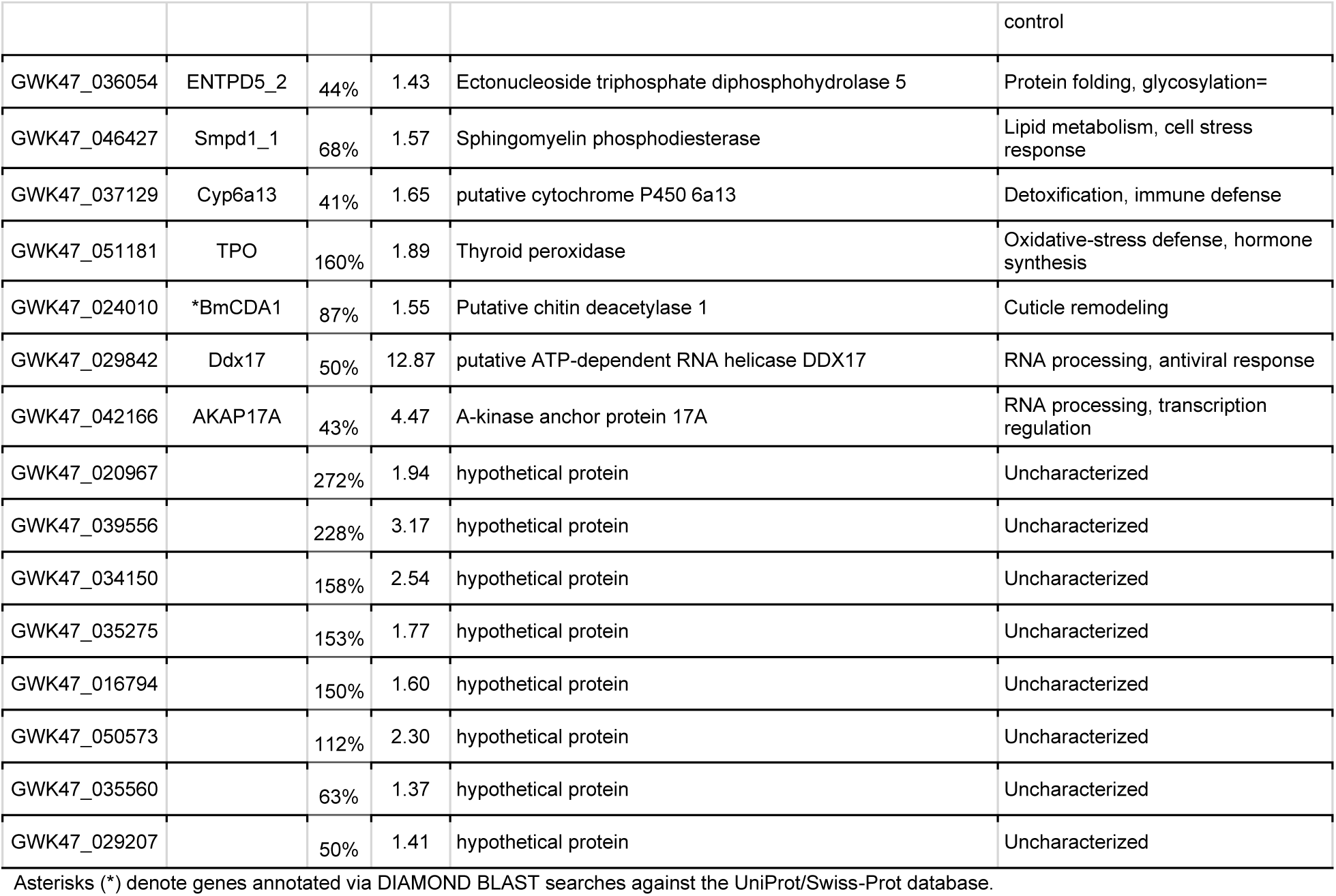
Top differentially expressed genes (|log_2_ fold change| > 0.5) distinguishing moderate vs. severe OA treatments after 88 days of exposure.

Genes with lower expression in moderate OA (n=87) were not significantly enriched for any terms. Annotated genes that were uniquely expressed in moderate OA with large effect size (Log_2_FC>1) included vesicular inhibitory amino acid transporter (slc32a1), and receptor-associated protein of the synapse (rpy-1) (Fig 3).

Severe acidification (pH 7.5) resulted in a larger transcriptional response, with 616 differentially expressed genes. Genes with higher expression in severe acidification (n=376) compared to control were highly enriched for the GO process mitochondrial translation (FDR=8.2e^−8^), and the Uniprot keywords respiratory chain (FDR=4.4e^−4^), electron transport (FDR=1.8e^−3^), and protein biosynthesis (FDR=0.023), and and the large effect-size genes (Log_2_FC≥1) that were uniquely expressed in severe OA included the lactosylceramide 4-alpha-galactosyltransferase (A4galt), a putative hydroxypyruvate isomerase (Hyi), Ficolin-2 (FCN2), ATP-dependent RNA helicase (Tdrd9), and putative nuclease HARBI1 (harbi1) (Figure 3). Genes with lower expression in severe acidification (n=240) were enriched for the GO processes cell adhesion (FDR=0.022), lipid transport (FDR=0.022), and response to endoplasmic reticulum stress (FDR=0.053), and annotated, large effect size genes (Log_2_FC>1) that were uniquely expressed at lower levels in severe OA included sodium-dependent phosphate transporter 1-A (slc20a1a), and protein trachealess (trh).

Forty-three genes were commonly differentially expressed in both OA treatments at 8-hr (blue points in Figure 3), 37 of which were annotated (Table S1). Genes that were expressed at higher levels in both OA treatments were enriched for the Uniprot keywords respiratory chain (FDR=0.038) and electron transport (FDR=0.079). No terms were enriched in genes with lower expression in both OA treatments.

### 3.4 Long term consequences: differential expression among OA treatments at Day 88

After 88 days of exposure, 186 genes were differentially expressed in severe OA (pH 7.5) compared to moderate OA (pH 7.8) (Table S5). A total of 80 genes expressed at higher levels in moderate OA compared to severe OA (blue points, Figure 4). While these genes were not significantly enriched for any biological processes, they represent possible mechanisms and markers of chronic OA tolerance in juvenile snow crab (see Table 3 for DEGs with L_2_FC>0.5, and Table S5 for all DEGs).

**Figure 4.**
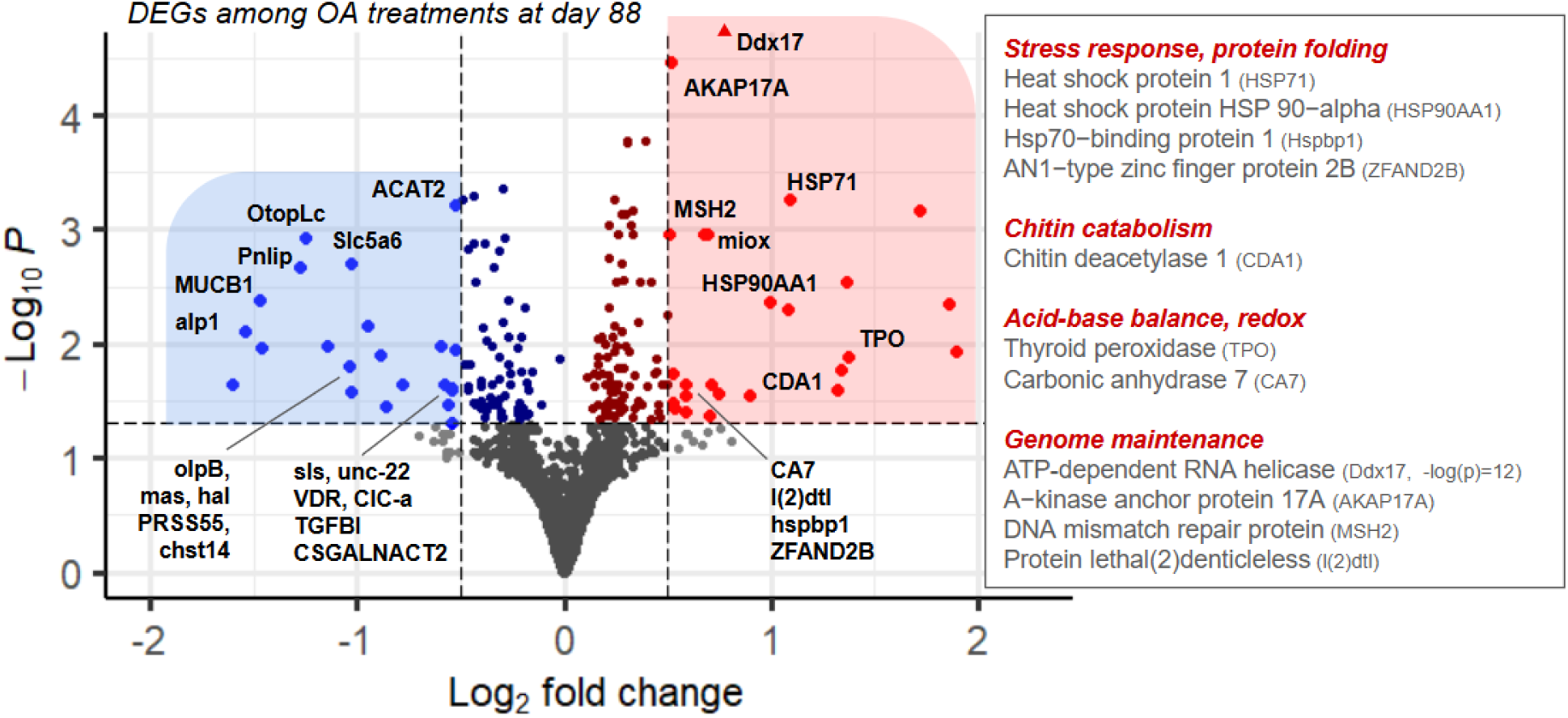
Volcano plot of differentially expressed genes (DEGs) between the moderate (pH 7.8) and severe (pH 7.5) ocean acidification (OA) treatments. Annotated genes meeting the significance thresholds of adjusted p < 0.05 and |log2 fold change| > 0.5 are in bright blue and red for genes with higher expression in moderate and severe OA, respectively. We highlight genes with higher expression in the severe OA treatment (red), which broadly represent stress-response, chitin catabolism, acid-base balance, and genome maintenance, which may underlie the higher long-term mortality observed in that treatment. See Table 3 for those less active in severe OA (blue points).

A total of 106 genes were expressed at higher levels in severe OA compared to moderate OA (red points, Figure 4), which were highly enriched for GO process mRNA processing (FDR=5.7e^−5^) and RNA splicing (FDR=9.2e^−5^). Genes that were highly expressed in severe OA (L_2_FC>0.5, or >1.4x) could be indicators of chronic OA stress (Table 2). Two annotated genes showed distinct expression patterns in long-term severe OA, and are therefore top candidates for OA stress biomarker development: carbonic anhydrase 7 (CA7) and protein lethal(2) denticleless (l(2)dtl) (Figure 5). Several additional genes were highly expressed in chronic severe OA exposure (*e.g.*, HSP90AA1, HSP71, Cyp6a13, Ddx17, AKAP17A) compared to chronic moderate OA, however they were also highly expressed in time-0 controls, indicating that they are likely sensitive to other factors and stressors (Power et al., 2023), and may not be reliable indicators of OA stress if measured alone.

**Figure 5.**
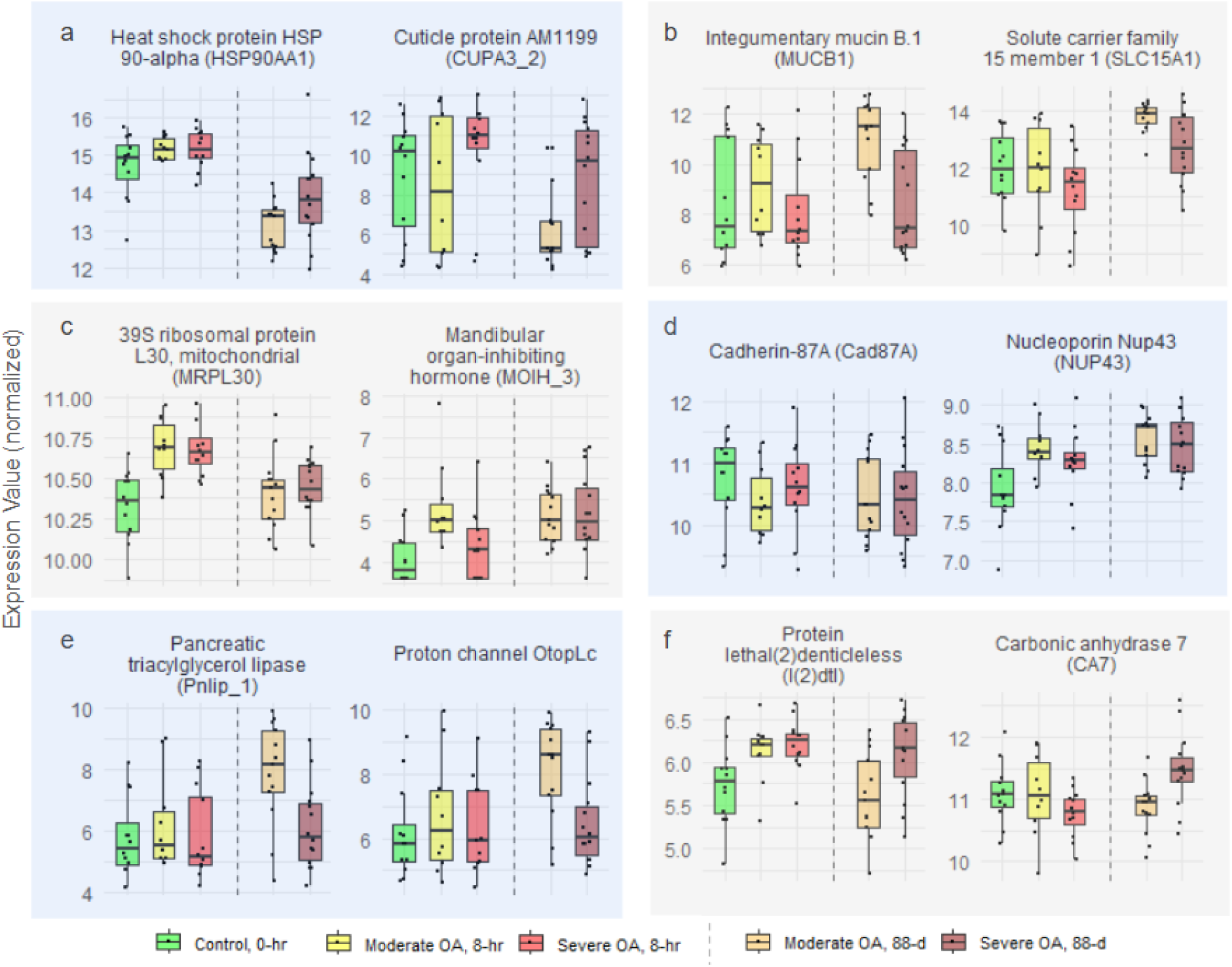
Expression patterns of select genes representing major axes of transcriptional variation and functional responses to ocean acidification (OA): (a) genes with high negative PC1 loadings, highly expressed in crabs likely experiencing OA stress; (b) genes with high positive PC1 loadings, highly expressed in crabs acclimated long-term to moderate OA; (c) genes differentially expressed in both OA treatments at 8 hr (MRPL30) or 88 days (MOIH); (d) genes that were differentially expressed in both OA treatments at 8 hr and 88 days; (e) genes uniquely or highly expressed in individuals acclimated to and tolerant of long-term moderate OA; and (f) genes highly expressed after 88 days of severe OA exposure, representing candidate biomarkers of chronic acidification stress. Boxplots show normalized gene counts without surrogate-variable (SV) correction. Dashed vertical lines separate sample days (Day 1 and Day 88).

Interactive boxplots of gene expression by treatment for all detected genes are available at: https://i0o16k-laura0h0spencer.shinyapps.io/snow_crab_expression_explorer/.

## 4. Discussion

Snow crab appear remarkably tolerant to ocean acidification compared with many other decapod crustaceans (Algayer et al., 2023; Long et al., 2023), even during the juvenile stage when other species show high sensitivity (e.g., red king crab: Long et al., 2013; Swiney et al., 2017). Long (2026) reported no detectable effect on juvenile growth or morphology during a full year of OA treatments. However, Long (2026) also revealed that snow crab tolerance has limits, with elevated molting-related mortality emerging after 250 days under severe acidification (pH 7.5), an effect that was not observed under moderate acidification (pH 7.8). In this discussion, we describe results from transcriptomic analyses conducted during the immediate acclimation period (8 h) and at an intermediate time point (88 d) that preceded mortality, to understand how snow crab tolerate OA, but why chronic exposure to severe acidification ultimately exceeds juveniles’ physiological capacity. Specifically, we describe (1) the major sources of gene expression variation across all treatments and time points, reflecting processes central to snow OA acclimation, (2) a more detailed description of the OA-responsive genes and processes that likely underpin short-term tolerance, and (3) transcriptional differences between moderate and severe OA at 88 days that reveal potential causes of mortality, and also identify candidate biomarkers distinguishing snow crab that are tolerating chronic OA and those that are experiencing chronic OA stress.

### 4.1 Largest signals of variation include cellular stress-response and cuticle maintenance genes

We conducted principal component analysis (PCA) of all 11,097 expressed genes after controlling for unknown surrogate variables (Figure 2), which revealed variation among treatments. The main axis of variability (PC1; Figure 2), which separated samples by treatment and exposure duration, was strongly influenced by heat shock protein expression (HSPs, Table S1). HSPs act as molecular chaperones that stabilize and refold misfolded proteins under stress (Roberts et al., 2010), and their induction is a hallmark of the cellular stress response in metazoans exposed to environmental stressors, including crustaceans under pH stress (Harms et al., 2014; Kumar et al., 2022; P. Li, 2017). During the initial acclimation period (8 h), expression of several HSPs was elevated in both OA treatments relative to the control. By 88-days, HSP expression in both OA treatments had subsided compared to initial levels. However, HSPs at day 88 were higher in the severe OA treatment (pH 7.5) compared to moderate OA (pH 7.8), suggesting that chronic pH 7.5 causes cellular stress requiring ongoing protein repair that does not occur, or occurs at lower levels, in pH 7.8. Together, these patterns indicate that HSP induction occurs both as an acute acclimation response and as a marker of sustained stress under long-term, severe acidification. Interestingly, the control group itself exhibited high HSP expression at 0-hr – higher than that observed in either OA treatment at day 88 – likely reflecting transient handling, transport, or general laboratory acclimation stress that subsided after 88 days (see also Power et al., 2023). Therefore, while HSPs are involved in OA acclimation, they are also more general stress-response mechanisms.

The top PC loadings also revealed clear treatment effects on genes involved in chitin metabolism and cuticle structure. Two genes coding for the cuticle protein AM1199 (CUPA3), a component of the chitin-based exoskeletal matrix (Liu et al., 2024), was highly and consistently expressed in severe OA crab during the short-term exposure (8 h, Figure 5), suggesting an early protective response aimed at stabilizing or reinforcing the cuticle. During the long-term exposure (88 d), AM1199 and other cuticle-related genes (putative CDA1, CU1A) remained elevated in severe OA relative to moderate OA. Consistent with sustained investment in cuticle maintenance or remodeling in OA, chitin-processing genes are associated with molting (Campli et al., 2024). For instance, expression of chitin deacetylase (CDA1), which modifies chitin structure by influencing Ca^2+^ and protein binding (Zhang et al., 2021), increases during molting and its disruption impairs molting in the Chinese mitten crab (X. Li, Chu, et al., 2021; X. Li, Diao, et al., 2021). While snow crab are able to maintain their exoskeleton properties on both the micro and macro scale (Algayer et al. 2023), the process of continuously building and maintaining cuticular tissues likely causes long-term energetic cost during chronic exposure, which may cause energetic challenges during molting that contribute to elevated mortality (Long et al., 2025). Some observed expression signals in snow crab may also reflect modifications to chitinous structures on the gill or gut as well as the exoskeleton (Novikov et al., 2023; Sarmiento et al., 2016; Zhang et al., 2021), an idea that warrants tissue-specific investigation.

It is notable that inter-individual variability was highest among snow crabs likely experiencing OA-stress, i.e. those in both the 8-hr OA treatments and the 88-day severe OA treatment (Figure 2). In contrast, inter-individual variability was lowest among red king crab juveniles experiencing the most severe OA treatment (Spencer et al. 2024). We posit that snow crab, as the more OA-tolerant species, is able to maintain higher expression across a more varied set of physiological processes (Rivera et al. 2023). Red king crab, as the less OA-tolerant species, may maintain a more suppressed metabolic and transcriptional state when OA-stressed. It would be interesting to test the hypothesis that inter-individual variability in expression profiles (high/low) reflects metabolic changes (acceleration/suppression) that predicts a species’ OA tolerance.

### 4.2 Short-term OA tolerance: acclimation mechanisms (8-hr response)

#### 4.2.1 Commonly affected processes & genes

Snow crab tolerance to ocean acidification may partly stem from their ability to activate key metabolic and protective pathways while minimizing unnecessary or costly ones. After short-term exposure (8 h), snow crabs in both OA treatments highly expressed genes involved in aerobic respiration and mitochondrial metabolism, with the response being far more pronounced under severe OA (pH 7.5). Approximately 30% of all genes highly expressed in severe OA were mitochondrial, including numerous ribosomal subunits, translation factors, respiratory chain components, and assembly proteins. Such responses suggest a strong cellular investment in maintaining mitochondrial function and increasing energy production capacity under acidified conditions. Enhanced mitochondrial performance likely underpins snow crab’s short-term tolerance to OA and may be essential for supporting protein repair and other energy-intensive defense and maintenance processes. Future studies could target metabolic genes (Table S2) across a wider range of pH levels to identify the threshold at which metabolic depression begins to occur in snow crab (Bednaršek et al., 2021; McElhany & Busch, 2024; Harms et al., 2014).

About one third of the genes differentially expressed under moderate OA at 8 h were also upregulated under severe OA, indicating a core set of genes that are likely essential for OA acclimation (Table S2). In addition to mitochondrial and metabolic genes, these included genes involved in β-oxidation, carbohydrate metabolism, protein processing, transcriptional regulation, and intracellular transport. One such gene, NUP43, encodes a nucleoporin and was elevated in both treatments and remained upregulated after 88 days (Figure 5). A related nucleoporin (NUP54) was identified as a hub gene during prolonged high-pH stress in shrimp (*Litopenaeus vannamei*; Huang et al., 2018), where its interaction with HSPA4 likely promoted mRNA export through the nuclear pore complex. NUP43 may serve a similar function in snow crab, helping maintain mRNA trafficking and protein synthesis under acidified conditions.

Genes expressed at lower levels in both short-term OA treatments were associated with structural maintenance and immune function. Several genes linked to cell adhesion and cytoskeletal organization (TGFBI, Cad87A, sas, rhpn2, MYO18A, CycG) were less abundant, including a gene coding for Cadherin 87A, which remained low after 88 days in both OA treatments (Figure 5). Two proclotting enzymes (PCE) were likewise reduced, consistent with short-term suppression of immune activity. It is possible that, rather than simple transcriptional downregulation, these patterns may reflect high metabolic or protein demand outpacing transcription rate. Because RNA-seq measures steady-state mRNA abundance, apparent decreases can result from reduced transcription, increased mRNA decay, or rapid turnover (Schwanhäusser et al., 2011). Future studies that sample more frequently during early acclimation could help distinguish transient mRNA fluctuations from true gene suppression.

#### 4.2.2 Dose-dependent OA effects: endocrine and osmotic regulation vs. damage mitigation

Differences between moderate and severe OA responses at 8 h indicate that snow crab tolerance operates along a continuum, where mild acidification elicits compensatory adjustments, and stronger acidification triggers cellular defense mechanisms. A gene in the crustacean hyperglycemic hormone (CHH) superfamily (Fanjul-Moles, 2006) showed markedly higher expression under moderate OA at 8 h (annotated as MOIH; Fig. 3). Although MOIH encodes the mandibular organ-inhibiting hormone, a neuropeptide implicated in suppressing vitellogenesis (Ding et al., 2023), similar transcripts were also elevated under transport stress in snow crab (Power et al., 2023), suggesting a broader stress-related function. Power et al. (2023) further proposed that these transcripts may represent other CHH-family members involved in regulating hemolymph glucose (Stoner, 2012; Webster et al., 2012), with downstream effects on metabolism and ion balance. Consistent with this interpretation, CHH-family neuropeptides are known regulators of ion and osmoregulation in decapods (Spanings-Pierrot et al., 2000; Chen et al., 2020). Indeed, we found that several genes highly expression in moderate OA are involved in osmotic and solute regulation, including SLC22A13, which facilitates organic ion and metabolite balance (Nigam, 2018), and SLC5A3, which imports compatible osmolytes to maintain cell volume and osmotic stability (Chauvin & Griswold, 2004). Given that crabs in this study were immature juveniles and years away from vitellogenesis, high levels of MOIH/CHH-like transcripts, which actually persisted after 88 days, likely reflects a central endocrine role in coordinating metabolic and osmotic homeostasis under moderate OA.

Genes elevated only under severe OA at 8 hrs were predominantly associated with cellular protection and damage mitigation, including TDRD9 and HARBI1, which function in transposon silencing and DNA repair (Gan et al., 2019; Kapitonov & Jurka, 2004), and FCN2, linked to innate immune activation and apoptosis (Jensen et al., 2007). Severe OA also elicited differential expression of heat shock proteins (as previously described), and multiple ion channels, including calcium- and potassium-permeable channels (e.g., TRPA1, Cacna1g, Ork1), which are known to participate in stress sensing/signaling and membrane stabilization. Together, these patterns indicate that moderate OA elicits low-cost regulatory compensation while severe OA shifts transcription toward cellular defense.

Notably absent from the short-term response to either OA treatment was transcriptional changes in most canonical ion-transport genes, including Na⁺/K⁺-ATPase and V-type H⁺-ATPase (V-ATPase). Although several components of these pathways were expressed, only TCIRG1, a putative V-ATPase subunit, showed treatment-dependent expression, with higher mRNA levels under moderate compared to severe OA. This limited response indicates that juvenile snow crabs do not rapidly encode ion-pumps during acute acidification, consistent with previous work showing that juvenile snow crabs hyporegulate magnesium only (Charmantier & Charmantier-Daures, 1995), and expectations for a species adapted to deeper, more stable conditions (Pane & Barry, 2007). Instead, differential expression of osmolyte transporters, particularly under more moderate acidification, suggests that snow crabs rely on hormonally mediated solute handling during immediate acclimation, likely resulting in partial conforming to ambient pH and pCO2. It is possible, however, that acid-base regulation is occurring with existing cellular machinery and therefore without transcriptional signal, or that it is spatially restricted to gill tissue and underrepresented in these whole-body transcriptomes. Monitoring hemolymph acid-base parameters during acclimation and targeted gill-specific expression studies will be required to fully resolve these mechanisms.

### 4.3 Molecular signatures of chronic OA exposure

After 88 days, crabs showed no outward signs of stress in survival, growth, or molt timing. However, the two OA treatments likely represent distinct positions along the stress-performance continuum. Crabs under moderate OA (pH 7.8) were probably functioning within their tolerance range, near the physiological optimum or entering the pejus zone, whereas those under severe OA (pH 7.5) were operating under suboptimal conditions. This interpretation aligns with the delayed but substantial mortality observed after ∼250 days in the severe treatment, which ultimately was appr. 40% higher than the moderate group (Long 2026). The transcriptional profiles captured at day 88 therefore provide an early glimpse into molecular processes distinguishing sustained tolerance from accumulating stress.

#### 4.3.1 Signatures of a crab tolerant of chronic, moderate OA

We identified dozens of genes with elevated expression after 88 days under moderate OA, which may contribute to snow crab’s long-term resilience to acidified conditions (Table 2 & Table S5). Many encode secreted or membrane-associated proteins, including integumentary mucin B.1 (MUCB1, Figure 5), serine protease 55 (PRSS55, also annotated to PROC, a vitamin K-dependent protein C), and sodium-dependent multivitamin transporter (Slc5a6). MUCB1 forms protective mucin coatings on epithelial surfaces (Probst et al., 1990), while PROC/PRSS55likely regulates blood coagulation to protect endothelial cells, and Slc5a6 mediates Na^+^-coupled vitamin uptake and is essential for gut mucosa maintenance in mice (Sabui et al., 2016), suggesting reinforced epithelial and mucosal defenses through enhanced secretion, proteolysis, and nutrient uptake. High expression of metabolic enzymes such as pancreatic triacylglycerol lipase (Pnlip, Figure 5) further points to increased lipid turnover supporting energetic and membrane demands (Lowe, 2002). Elevated expression of ion-transport genes, notably the proton channel OtopLc (Figure 5) and the chloride channel ClC-a, implies differences in ion regulation (Cabrero et al., 2014; W. W. Chang et al., 2021; Wheatly & Henry, 1992). Although we did not detect differences in major ion exchangers (Na⁺/K⁺-ATPase or V-type H⁺-ATPase), these treatment-dependent differences in ion channels raise the possibility that acid-base regulation occurs during chronic OA exposure in snow crab. However, the absence of an ambient day-88 control prevents direct comparison with the long-term acid–base regulation observed in Tanner crab (Meseck et al., 2016). Several immune-regulatory genes were also upregulated, including masquerade (mas) which potentially moderate inflammation levels, cell adhesion, and epithelial integrity (Murugasu-Oei et al., 1995), and a gene similar to cybc1 (GWK47_039981), which promotes reactive oxygen species production for pathogen defense (Arnadottir et al., 2018). High expression of adhesion, and structural genes, including TGFBI, OLPB₂, Titin, and two Twitchin paralogs, suggest coordinated maintenance of muscle function (Ayme-Southgate et al., 1991; Murugasu-Oei et al., 1995).

Collectively, the dominant transcriptional signals under long-term exposure to moderate OA indicate enhanced secretion of protective proteins and enzymes, strengthened epithelial barriers, stabilized ion gradients, and maintained immune and structural function compared to those in severe OA.

#### 4.3.2 Signatures and candidate biomarkers of chronic OA stress

Genes highly expressed after 88 days under severe OA provide mechanistic insight into chronic acidification stress that, as shown by Long (2026), ultimately leads to elevated molt-associated mortality in juvenile snow crabs. Broadly, genes upregulated in severe OA relative to moderate OA were involved in protein folding, genome maintenance, transcriptional regulation, pH homeostasis, and immune and metabolic control (Table 2). Many of these same processes were elevated in severe OA during the initial acclimation phase, indicating that exposure to pH 7.5 constitutes a stressor despite long-term exposure.

Genes involved in chaperone-mediated refolding (*e.g.*, HSP71, HSP90AA1, Hspbp1) and proteasomal degradation (*e.g.*, ZFAND2B, l(2)dtl) were among the most highly expressed overall. HSP synthesis, and protein synthesis more broadly, is energetically costly (Feder & Hofmann, 1999). However, Long (2026) saw no differences in growth among treatments over a one-year exposure, implying that energetic demands were met when crabs were not molting.

Instead, the molt-associated mortality observed in severe OA could reflect the combined physiological strain of molting and acidification exceeding the crab’s capacity for cellular repair. Indeed, HSP expression varies among molt stages in other decapods (Cesar & Yang, 2007; E. S. Chang, 2005; López-Cerón, 2019; Spees et al., 2003). Despite their strong responses, HSPs are not ideal standalone biomarkers of chronic OA exposure because they are broadly inducible by diverse stressors (*e.g.*, as the name suggests, they were originally described in response to high temperature stress) (De Maio et al., 2012; Kumar et al., 2022). In our study, HSP expression was higher in control crabs sampled on Day 1 than in OA-exposed crabs at Day 88, potentially reflecting handling stress, or exposure to more variable environmental conditions prior to the experiment (among other possible stressors) (Power et al., 2023). We therefore do not recommend HSPs as standalone indicators of OA stress.

Carbonic anhydrase 7 (CA7 or CAH7; GWK47_031192) was the most distinctively upregulated gene in crabs exposed to long-term severe OA (Figure 5) and represents our top candidate for biomarker development. In crustaceans, carbonic anhydrases (CAs) are highly active in gill tissue, where they support acid-base regulation, CO_2_ excretion, and provide bicarbonate for CaCO_3_ formation during calcification (Henry, 1988; Le Roy et al., 2014; McMahon et al., 1984). Although CA7 has not been characterized in crustaceans, studies in vertebrates identify it as a cytosolic isoform that catalyzes the reversible hydration of CO_2_ to bicarbonate and protons (Aspatwar et al., 2022). Because of their central roles in ion regulation and biomineralization, CAs have been widely proposed as biomarkers of environmental stress in calcifying animals (Zebral et al., 2019). Numerous studies in corals and molluscs have reported altered CA transcription and enzyme activity under acidified conditions, most commonly showing suppression (reviewed in (Zebral et al., 2019)). In contrast, CA7 expression in snow crab increased, and only after prolonged exposure to severe acidification rather than during the initial 8-h response. This delayed activation suggests that CA7 reflects a shift from an immediate OA response to chronic acclimation, making it a promising molecular indicator of sustained OA exposure. Future work should identify the tissue(s) in which CA7 expression is most responsive to OA, as the present analysis used whole-body homogenates.

A second notable gene, lethal (2) denticleless (l(2)dtl), was uniquely expressed in both short-term OA treatments and the long-term severe OA treatment (Figure 5). Conserved across taxa, l(2)dtl regulates DNA replication and repair via ubiquitin-mediated degradation pathways (Sloan et al., 2012). Its high expression under exposure to OA may reflect enhanced DNA damage surveillance or adaptation to replication stress, making it a potentially sensitive indicator of genotoxic stress.

## 5. Conclusion

Snow crabs show a remarkable ability to maintain growth under ocean acidification (Long 2026). This study described the flexible transcriptional responses that appear to buffer cellular stress.

Even after months of exposure to low pH, juveniles sustained heightened expression of genes involved in protein repair and cuticle maintenance, with no detectable effects on growth or molting (Long 2026). These results suggest that snow crabs tolerate acidified conditions through active molecular repair. A similar pattern has been observed in postlarval American lobster, where strong activation of stress and immune pathways maintained performance under OA (Niemisto et al., 2021). In contrast, species such as the Antarctic pteropod (*Limacina helicina antarctica*, Johnson and Hofmann 2017) and the Chinese mitten crab (*Eriocheir sinensis*, Luo et al. 2021) exhibit transcriptional suppression or trade-offs between defense and reproduction, consistent with reduced plasticity and lower resilience to acidification.

The eventual molt-associated mortality observed under chronic severe acidification indicates that snow crabs’ tolerance has limits. Genes such as carbonic anhydrase 7 (CA7) and l(2)dtl have potential to be used to detect whether a snow crab is experiencing OA stress, which could be a useful tool for monitoring Bering Sea populations (Litzow et al., 2025; Szuwalski et al., 2021). Future work should focus on tissue-specific expression with finer temporal resolution to better understand the timing and location of OA-responsive genes. Comparative analyses with the closely related, OA-sensitive Tanner crab will be especially valuable for identifying the molecular mechanisms underlying differences in resilience among North Pacific crab species. These and other molecular analyses would benefit substantially from improved annotation of genes and other regulatory features in the snow crab genome assembly.

## Acknowledgements

Thank you to NOAA Eastern Bering Sea Bottom Trawl Survey team for their collection and transport efforts, Giles Goetz, Mia Nahom, and the rest of the Sedna team at the NWFSC for computing capabilities and support, and to Cody Szuwalski for the helpful comments.

## Data and resource availability

Raw sequence data will be available on NCBI SRA upon publication; additional data and code is available on GitHub, https://github.com/laurahspencer/snow-crab_RNASeq-2022.

