## Supplementary for "Short-term mechanisms, long-term consequences: molecular effects of ocean acidification on juvenile snow crab"

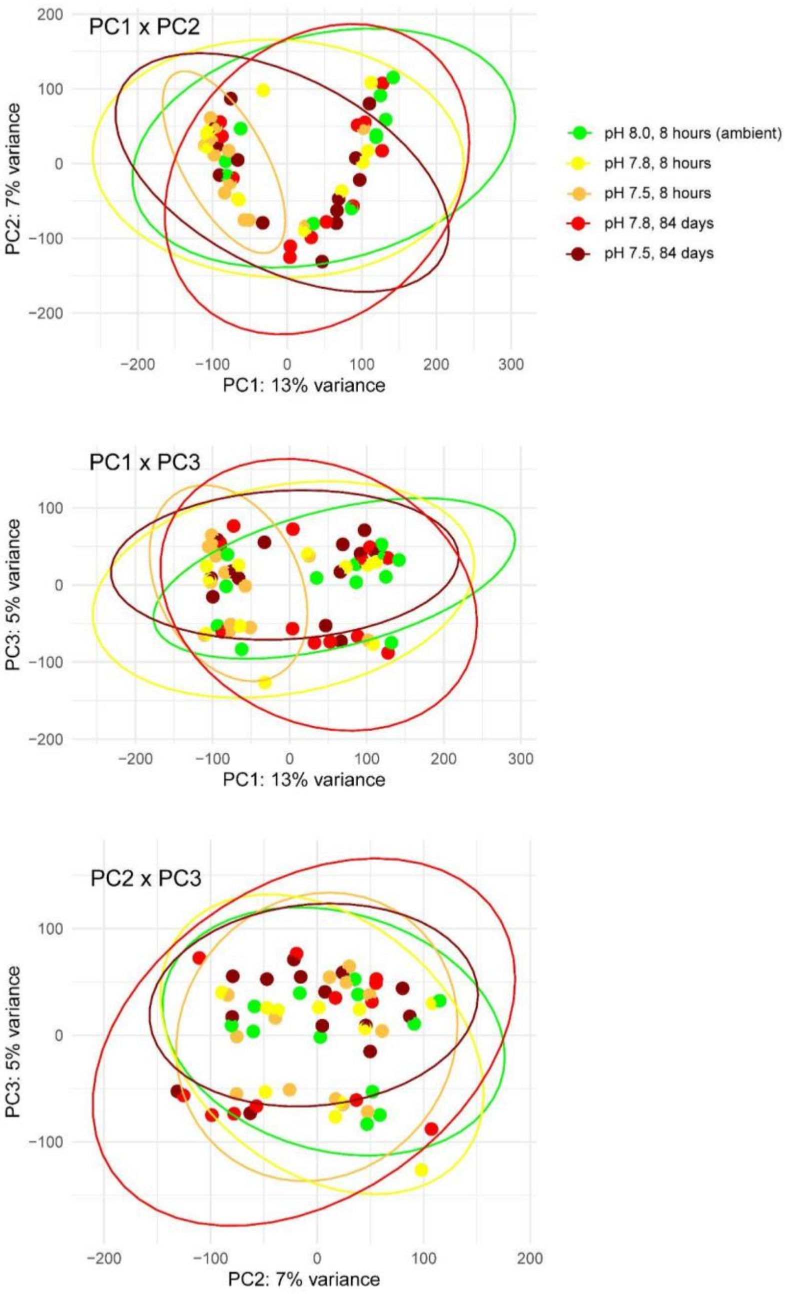

Figure S1: Principal component analysis biplots (PC1-PC3) without including surrogate variable analysis to control for unknown latent effects. The horseshoe pattern in PC1xPC2 and clustering by factors not related to treatment in biplots along PC1 and PC3 demonstrate the expression heterogeneity of technical or biological variables that obfuscates acclimatory response to acidification.

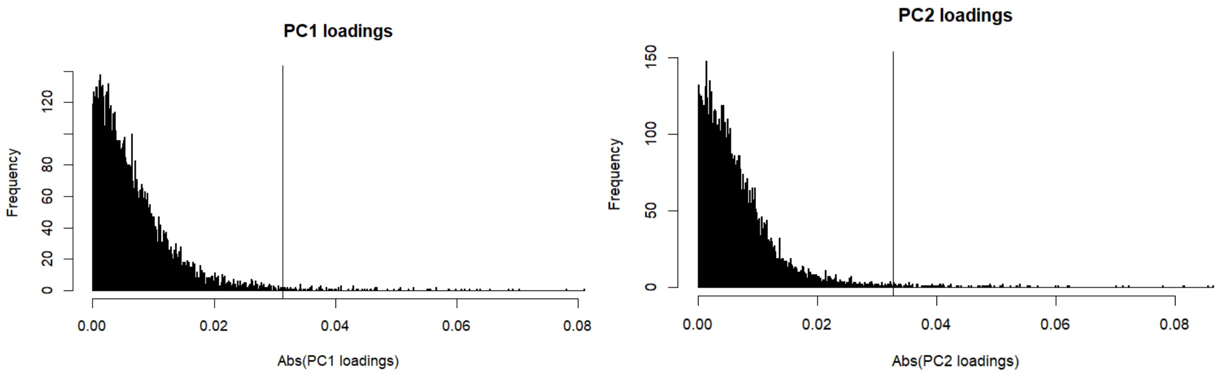

**Figure S2**: Histograms of PC1 & PC2 loadings (absolute values) from the VST- and SVA-controlled count matrix, the top 1% of which were >0.032 (to the right of vertical lines).

**Table S1**: Highlighted list of annotated genes that were differentially expressed in both moderate (pH 7.8) and severe (pH 7.5) OA after 8-hr exposure. $Log_2$ Fold Change ($L_2FC$) and -Log(P-adjusted) are compared to control crab held in ambient conditions (pH 8.0) at time-0. For all genes, the relative expression compared to control (higher, lower) was consistent in both OA treatments. **Indicates genes that were also differentially expressed after 88-days in both treatments.

| Gene Name | Gene ID | Mod. OA Δ% | Mod. OA -Log(P) | Sev. OA Δ% | Sev. OA -Log(P) | Protein | Biological Processes |
|---|---|---|---|---|---|---|---|
| **Higher Expression in both OA treatments at 8 h** | | | | | | | |
| Ndufs2 | GWK47_042301 | 22% | 1.45 | 25% | 2.34 | NADH dehydrogenase [ubiquinone] iron-sulfur protein 2, mitochondrial | Mitochondrial respiration, energy metabolism |
| Ndufs3 | GWK47_035555 | 19% | 1.42 | 28% | 3.15 | NADH dehydrogenase [ubiquinone] iron-sulfur protein 3, mitochondrial | Mitochondrial respiration, energy metabolism, redox |
| NDUBA | GWK47_048997 | 20% | 1.36 | 27% | 2.61 | NADH dehydrogenase [ubiquinone] 1 beta subcomplex subunit 10 | Mitochondrial respiration, energy metabolism |
| MRPS22 | GWK47_024976 | 24% | 1.98 | 21% | 2.32 | 28S ribosomal protein S22, mitochondrial | Mitochondrial translation, energy metabolism |
| mRpL35 | GWK47_003027 | 20% | 1.43 | 20% | 1.96 | 39S ribosomal protein L35, mitochondrial | Mitochondrial translation, energy metabolism |
| MRPL30 | GWK47_027395 | 18% | 1.36 | 27% | 2.97 | 39S ribosomal protein L30, mitochondrial | Mitochondrial translation, energy metabolism |
| UGT2B15_3 | GWK47_007253 | 117% | 1.52 | 7% | 1.31 | UDP-glucuronosyltransferase 2B15 | Detoxification, xenobiotic metabolism |
| 4CL_0 | GWK47_015047 | 110% | 1.87 | 52% | 1.36 | 4-coumarate--CoA ligase | Energy metabolism |
| Echs1 | GWK47_021840 | 21% | 1.60 | 17% | 1.65 | Enoyl-CoA hydratase, mitochondrial | Lipid metabolism, β-oxidation |
| miox | GWK47_050003 | 40% | 1.31 | 44% | 2.07 | Inositol oxygenase | Carbohydrate metabolism |
| emc2-b | GWK47_005824 | 23% | 1.93 | 13% | 1.32 | ER membrane protein complex subunit 2-B | Protein processing, ER membrane insertion |
| Xpnpep1_2 | GWK47_044723 | 11% | 1.41 | 9% | 1.38 | Xaa-Pro aminopeptidase 1 | Protein turnover, stress response |
| MED27 | GWK47_029699 | 17% | 1.36 | 16% | 1.79 | Mediator of RNA polymerase II transcription subunit 27 | Transcriptional regulation (RNA polymerase II) |
| prmt6 | GWK47_007501 | 24% | 1.87 | 26% | 2.79 | Protein arginine N-methyltransferase 6 | Epigenetic and transcriptional regulation |
| Srsf7 | GWK47_020456 | 16% | 1.36 | 16% | 2.06 | Serine/arginine-rich splicing factor 7 | RNA splicing, mRNA processing |
| U2af50 | GWK47_001080 | 15% | 1.49 | 13% | 1.56 | Splicing factor U2AF subunit | RNA splicing, mRNA processing |
| Tsnax | GWK47_041448 | 14% | 1.41 | 16% | 2.22 | Translin-associated protein X | RNA silencing, reproductive process |
| Myo5a_0 | GWK47_011520 | 61% | 2.46 | 27% | 1.44 | Unconventional myosin-Va | Cytoskeletal transport, vesicle trafficking |
| **NUP43 | GWK47_010369 | 27% | 1.41 | 21% | 1.45 | Nucleoporin Nup43 | Nucleocytoplasmic transport, cell cycle regulation |
| **Lower Expression in both OA treatments at 8 h** | | | | | | | |
| oct-1_1 | GWK47_003819 | -71% | 1.93 | -61% | 1.90 | Organic cation transporter 1 | Ion transport, acid–base homeostasis |
| Gstt4 | GWK47_034834 | -41% | 2.38 | -23% | 1.32 | Glutathione S-transferase theta-4 | Detoxification, oxidative stress response |
| RSPRY1_1 | GWK47_007365 | -17% | 1.34 | -16% | 1.62 | RING finger and SPRY domain-containing protein 1 | Protein degradation, turnover |
| USP19 | GWK47_030223 | -15% | 1.38 | -14% | 1.55 | Ubiquitin carboxyl-terminal hydrolase 19 | Protein quality control, stress response |
| Phm | GWK47_018134 | -46% | 1.84 | -34% | 1.63 | Peptidylglycine alpha-hydroxylating monooxygenase | Peptide processing, reproduction |

| | | | | | | | |
|---|---|---|---|---|---|---|---|
| PCE_0 | GWK47_027739 | -43% | 1.63 | -41% | 2.17 | Proclotting enzyme | Immune, coagulation response |
| PCE_2 | GWK47_051844 | -35% | 1.37 | -43% | 2.64 | Proclotting enzyme | Immune, coagulation response |
| VPS13C_1 | GWK47_005657 | -27% | 1.31 | -30% | 2.10 | Vacuolar protein sorting-associated protein 13C | Lipid transport, organelle organization (mitochondrial) |
| PDXK | GWK47_032265 | -38% | 1.86 | -24% | 1.33 | Pyridoxal kinase | Cofactor metabolism, cellular metabolism |
| TGFBI | GWK47_026454 | -37% | 1.60 | -31% | 1.68 | Transforming growth factor-beta-induced protein ig-h3 | Cell adhesion, structural organization |
| **Cad87A | GWK47_018958 | -23% | 1.86 | -19% | 2.02 | Cadherin-87A | Cell adhesion, signaling |
| sas_3 | GWK47_043552 | -30% | 1.31 | -36% | 2.34 | putative epidermal cell surface receptor | Cell signaling, membrane receptor activity |
| rhpn2 | GWK47_028156 | -22% | 1.58 | -17% | 1.44 | Rhophilin-2 | Cytoskeletal signaling |
| MYO18A | GWK47_044957 | -18% | 1.52 | -15% | 1.45 | Unconventional myosin-XVIIIa | Cytoskeletal transport, cell organization |
| CycG | GWK47_003032 | -23% | 1.41 | -24% | 2.11 | Cyclin G | Cell cycle regulation, growth control |
| A1CF | GWK47_037491 | -21% | 1.36 | -20% | 1.78 | APOBEC1 complementation factor | Reproduction, RNA editing |
| DOP1A | GWK47_009521 | -15% | 1.61 | -15% | 2.27 | Protein dopey-1 | Vesicle trafficking, protein transport |

### Descriptions of Tables S2-S6

*Please also see Google Spreadsheet for Tables S2-S6 (in addition to .csv files).*

**Table S2**: Genes with top loadings along PC1 and PC2.

**Table S3**: Full list of DEGs after 8 hr OA exposure.

**Table S4**: Persistent DEGs, those from 8 hr that were also DEGs at 88 days.

**Table S5**: Lists of DEGs among OA treatments at 88 days.

**Table S6**: Enriched biological processes.

### Short term acclimation mechanisms that persisted at day 88

We did not generate expression data for the ambient treatment at day 88, so cannot fully separate OA effects from other potential influences of long-term laboratory exposure (e.g., diet, tank environment). To focus on genes most consistently associated with OA, we identified differentially expressed genes (DEGs) at 8 h that remained differentially expressed at 88 days (Figure S3, Table S4).

In moderate OA, 18 DEGs identified at 8-hr persisted at 88 d (Table S4), which were enriched for RNA binding (GO:0003723) and included several involved in transcription and splicing, proteolysis, retinol metabolism, cell adhesion/structure, and possibly circadian rhythm (Mandibular organ-inhibiting hormone, see Fig 3).

In the severe OA treatment, 59 DEGs identified at 8-hr persisted at 88 d (Table S4), which were not significant enriched for any processes, but broadly reflect genes involved in mitochondrial energy metabolism, detoxification, and defense against oxidative or hypoxic stress (HSP22, CYP2L1, SIRT4, CBS, Gstt4), cell division and transcriptional control (e.g. cdk1, Mafg, Usp21), cell adhesion and cytoskeleton dynamics (TTN, Actr3, Cad87), and hormone and growth factor signaling (FLT4, LHCGR, Tdrd9). These “persistent” DEGs suggest that severe OA induces long-lasting shifts in stress response, signaling, and cellular regulation (Table X).

Two genes were found to persist in both moderate and severe OA treatments: cad87A which codes for a cadherin, and nup43 which codes for a nucleoporin, and which were expressed at lower and higher levels, respectively, in OA treatments compared to control (Figure S4).

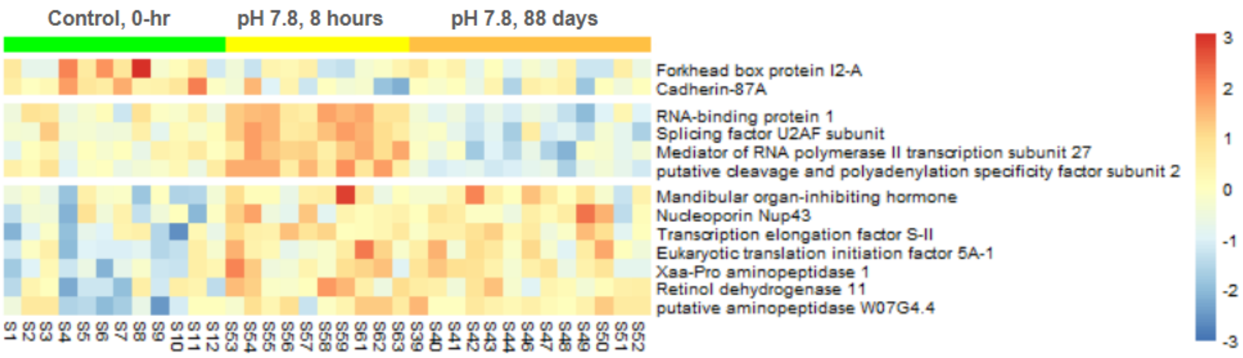

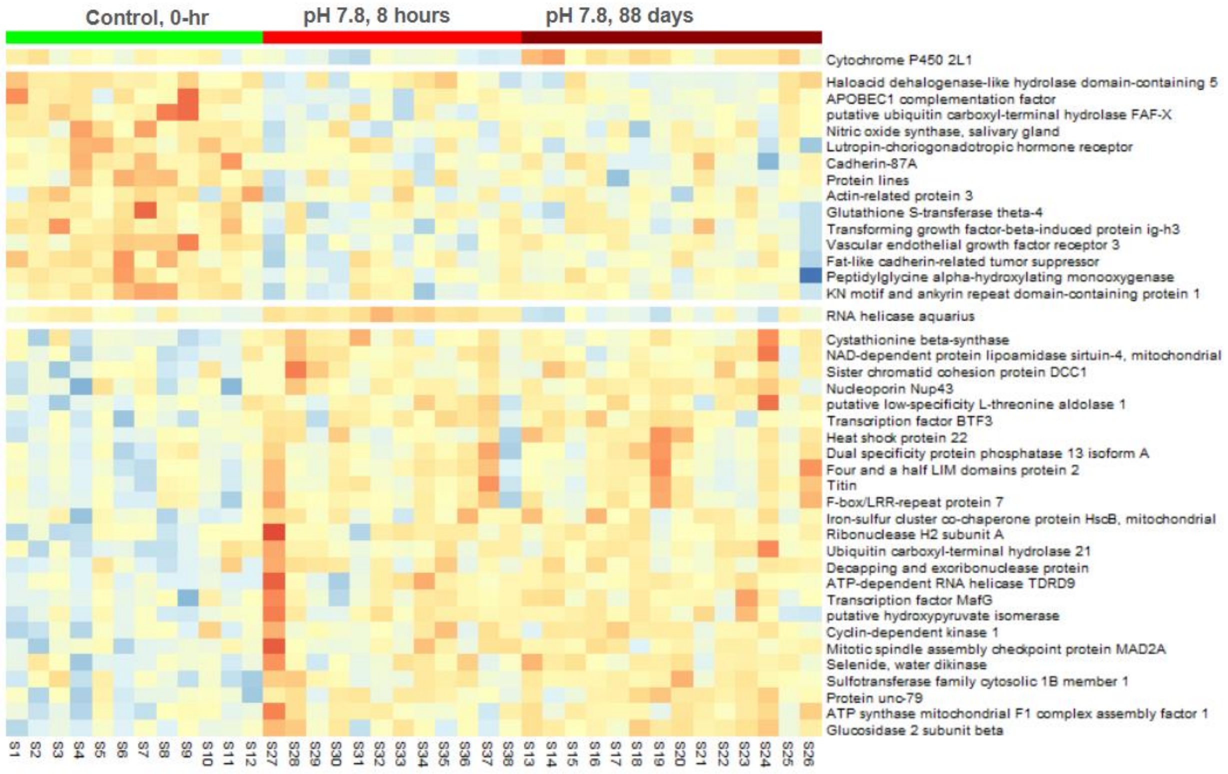

**Figure S3**: Heatmaps of persistent DEGs, defined as genes differentially expressed at both 8 h and 88 d. Results are shown for moderate OA (pH 7.8, top) and severe OA (pH 7.5, bottom) relative to control conditions measured at time 0. Colors indicate relative expression (z-scores), with red = higher and blue = lower expression, columns represent individual crabs, and rows represent individual genes, which are grouped by their expression pattern. Two genes were persistent DEGs in both OA treatments: cadherin-87A (cad87a) and nucleoporin (nup43), which were expressed at lower and higher levels, respectively, in OA compared to control.
